# Drug repositioning with gender perspective focused on Adverse Drug Reactions

**DOI:** 10.1101/2022.07.22.501091

**Authors:** Belén Otero Carrasco, Aurora Pérez Pérez, Ernestina Menasalvas Ruiz, Juan Pedro Caraça-Valente Hernández, Lucía Prieto Santamaría, Alejandro Rodríguez-González

## Abstract

Drug repositioning is a novel, useful, and crucial technique to find new uses for existing drugs. In this field of study, when the clinical trials necessary to obtain successful drug repositioning have been carried out, the female gender has not been given much consideration. Thus far, the participation of women in clinical trials has been very limited. There were several argued reasons to exclude them from trials, like the likelihood of pregnancy or sudden hormonal changes. This has meant that for a long time the adverse effects of a drug on women were unknown. Scientifically, it was known that due to the biological processes of pharmacokinetics and pharmacodynamics, the response to drugs was not the same in both genders, but despite this evidence, there is still no difference in the dosage or form of using a drug between men and women. In this study, we made a preliminary analysis where the main goal is to investigate gender differences within the drug repositioning field through the adverse effects produced by such treatments. A special section on specific cases of drug repositioning in rare diseases will also be considered to carry out the same verification previously mentioned in the text.

## I. Introduction

Historically, the omission of women from clinical trials has been based on the idea that the pharmacokinetics and the pharmacodynamics of a drug can be influenced by menstrual cycle phases, hormonal fluctuations, the use of oral contraceptives, and hormonal therapy, and life events such as pregnancy and lactation. Consequently, a considerable number of the drugs marketed over the years have been exclusively clinically tested on men [1].

In 1993, the FDA (Food and Drug Administration), through the Guideline for the Study and Evaluation of Gender Differences in the Clinical Evaluation of Drugs, called for the participation of women in clinical trials and recommended men and women be analyzed separately when it comes to drug responses [2] [3].

In 2001, the Institute of Medicine (IOM) of the US National Academy of Sciences published a report called “Exploring the Biological Contributions to Human Health – Does Sex Matter?”. The report emphasized that being male or female affects health, and how gender should be considered when designing or analyzing studies at all levels of biomedical research. The conclusion of the IOM report is crucial to keep in mind the potential differences between the genders while developing medicines [4].

Despite this, predominantly, women are still under-represented in clinical trials. Consequently, healthcare providers are often left to estimate the appropriate dose and interval for treatment or whether it is safe to use it during pregnancy without the necessary knowledge [5].

Several review studies have shown that the presence of women in clinical trials remains low. One of them shows that out of 132 clinical trials reviewed, only 40 of them met the inclusion criteria for women[6]. Another paper focused on the drug etoricoxib, which has been found to interact with female hormone therapy; despite this relationship, in the phase I of the study only 30% of the sample population were women [7].

Furthermore, males and females react to drugs divergently based on biological evidence focused on pharmacokinetics and pharmacodynamics. Within pharmacokinetics, there are gender differences in drug absorption (Gastric enzymes, Transporter proteins, Enterohepatic and renal handling of drugs or metabolites), distribution (Body fat composition, Cardiac output), metabolism (Hepatic metabolism), and elimination (Clearance from the body is slower in women) [1]. Of the above-listed differences, the one that most influences the women’s and men’s pharmacokinetics is metabolism [8].

In relation to pharmacodynamics, there are three main pharmacological factors that are affected to a greater extent in women. These factors are alteration in receptor number, alteration in receptor binding, and alteration in signal transduction pathway following receptor binding [5].

These factors affect the number of side effects produced by drugs, which is commonly higher in women than in men [1]. While these differences exist, most drugs do not provide sex-specific dosing instructions [9]. When women consistently experience less therapeutic effects or more adverse effects, a change in dosing regimen may be necessary or assume that the drug is not suitable for treating this pathology in women [10].

The manuscript has been organized as follows: Section 2 contains the related works on the exposed topics, Section 3 explains the methods used for the analysis, Section 4 presents the obtained results in the study, Section 4 details the discussion, and Section 5 presents the conclusions and future works.

## II. RELATED WORK

Drug repositioning from a gender perspective is a poorly explored field of study. There is very scarce scientific literature that considers both aspects together. However, drug repositioning and the gender perspective separately are two topics of great interest at present.

Several studies have shown that drug repositioning (DR) is a valuable and essential tool to find new uses for existing medicines [11]. It presents notable advantages against *de novo* drug discovery, as it can evade enormous costs, risks, and time. On the other hand, as has been previously explained, gender perspective has been poorly considered in *de novo* drug development, and for this reason, its consideration in drug repositioning is of utmost importance.

Relevant research based on this computational mechanism of drug repurposing has led to historically landmark discoveries. A few clear examples are thalidomide, sildenafil, or duloxetine, which were serendipitously identified to be efficient treatments for new indications [12].

The use of the drug repositioning process has become more common over time thanks to advancements in computer-based approaches, especially in artificial intelligence. A wide variety of artificial intelligence techniques and machine /deep learning algorithms have been used to develop new hypotheses about computational drug repositioning where techniques such as logistic regression [13], support vector machine [14] and neural networks [15], among others, have stood out.

A great example of a computer-based approach to generating drug repositioning hypotheses can be seen in this paper [16], that has established a methodological pipeline that can generate new drug repositioning hypotheses by integrating biomedical knowledge. In this line, [17] shows an approach using a data-driven approach to determine several potential drugs that could be repositioned for COVID-19, showing the importance and the potential impact of these techniques.

In this context, drug repositioning, and especially those approaches based on data-driven techniques are of extreme importance to find therapeutic options for diseases. This is something quite important for example in rare diseases, where typically the investment to find new drugs is much lower than in other pathologies, and where these approaches can help to find treatments for these diseases. Rare diseases usually pose more challenges to drug repositioning because there is a scarcity of data. Nevertheless, computational approaches have been developed based mainly on similarities between omics data and biochemical features of diseases [18].

Another important aspect that has been presented and is part of the core of this work is the gender perspective. Previous studies based upon gender differences in drug discovery show interesting results, which demonstrate that gender differences are present, and the importance of taking them into account.

Using data compiled from more than 500,000 patients collected throughout the 1982-97 period, Martin et al. [19] determined how the women over the age of 19 had a 43 to 69% greater likelihood of recording an Adverse Drug Reaction (ADR). Another study, which considered the 668 most frequently used drugs as treatments in the USA, found that 307 (46%) reported significant sex differences in ADRs [20].

FDA evaluation of 300 new drug applications between 1994 and 2000 [21] indicated that 31% of studies showed a possible sex effect based on pharmacokinetic sex differences greater than 20%. In this same study, despite obtaining differences over than 40% between men and women in pharmacokinetic measures, no gender-related drug dosing was recommended.

Across all drug classes, women have nearly double the risk of experiencing adverse drug reactions and are more likely to be treated in a hospital due to an adverse drug reaction [22].

Additionally, women are more likely than men to be on two or more medications at the same time (polypharmacy) and use more medications per year (5.0 vs. 3.7, respectively), which could contribute to an increase in adverse drug reactions in females [23] and puts a greater emphasis on sex-aware dosing. Women reached their maximum prevalence of ADRs earlier in the development process than men (30–39 years of age vs 50–59) [19].

The study of drug repositioning and the gender perspective, together, can lead to very interesting and important results for scientific research.

Recent evidence suggests that the pharmacokinetic differences between the genders have not been considered when evaluating the repositioned drugs proposed as potential treatments for COVID 19. It was determined therefore to be of utmost importance to advocate for a more gender-sensitive research initiative to develop more effective and safer therapies for women and men. [24].

## III. Methods

In this section, we proceed to explain the acquisition and integration of all relevant information for the two case studies considered: non-rare diseases and rare diseases.

In our research, the scientific literature has been queried for successful cases of drug repositioning.

These cases of drug repositioning are triplets composed of the drug name, the disease for which the drug was indicated originally, and the new indication for which it was repositioned (Fig.1).

**Fig. 1.**
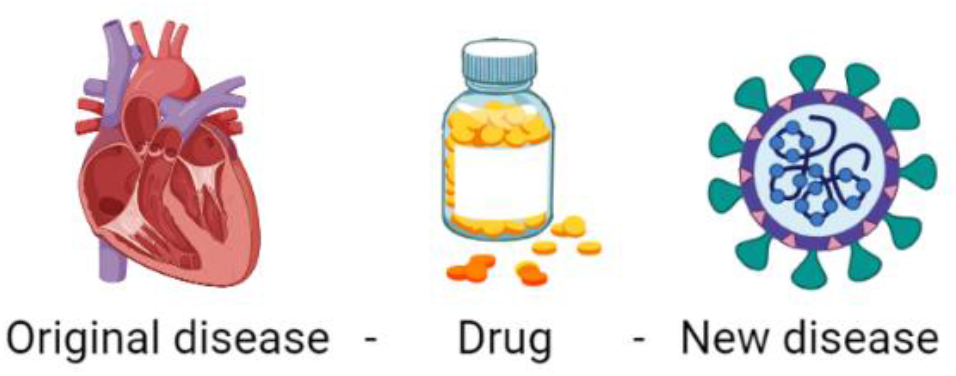
Description of the components of a drug repositioning triplet.

For non-rare diseases the data sources were predominantly three articles: Xue *et al* [25], Jarada *et al* [26], and Li and Jones [27]. This selection has been made based on a recent study where the selection of successful drug repositioning cases was further explored [16].

Focusing on drug repurposing for rare diseases, the triplets considered in this study have been mainly obtained from two articles: Scherman and Fetro [28] and Dhir et al [29]. For some of these successful repositioning cases, we have obtained literature explaining the process by which drug reuse occurred in that particular disease. This is the case, for example, for Dengue [30], Zika [31], Leishmaniasis [32], or Leprosy [33]. Many of the successful triplets found for rare diseases have been used to treat parasitic diseases.

We used DISNET [34] to obtain the necessary information about the drugs and diseases. So, we discarded the triplets whose diseases were not present in DISNET. DISNET (https://disnet.ctb.upm.es/) is a platform that contains biological information of a set of diseases and their relationship with several features, including phenotypic characteristics and drugs associated with the diseases. The information gathered consisted of the diseases’
s Concept Unique Identifier (CUI) and the drugs’ ChEMBL ID.

The CUI [35] was originally obtained from the Unified Medical Language System (UMLS) for each of the diseases involved in the collected cases of successful drug repositioning. Then, only those unique cases were selected from the different data sources mentioned above. This created the final dataset for the continuation of this study. As for the drugs, the ChEMBL ID [36] that is generated by the European EMBL Bioinformatics Institute (EMBL-EBI) was obtained to identify each of the drugs with a unique ID.

From this point onwards, the percentage value of ADRs for males and females for each of the drugs presented was obtained. This information was gathered through the VigiAccessTM website [37], which has been designed to provide greater transparency in pharmacovigilance and uses available data on possible side effects that have been reported to the World Health Organization Programme for International Drug Monitoring (WHO PIDM) [38].

In summary, for each successful triplet considered, we obtained the ADRs value by gender for the drug involved, and the global prevalence value for both diseases. After this data preprocessing, we applied the same process to both case studies, non-rare and rare diseases, as described below.

The first step was to perform a statistical analysis to determine whether there were differences between men and women in terms of ADRs. For this purpose, the Mann-Whitney U test was used to make the comparison between the two genders since the data studied did not follow a Gaussian distribution. To reach this conclusion, the relevant tests were carried out, the Levene Test for homogeneity of variances and, the Lilliefors normality test.

Then, the second step in our study was to carry out an analysis on the distribution of ADRs by gender also considering the diseases prevalence by gender. Therefore, it was necessary to acquire global prevalence values for each of the diseases (by gender) from the existing cases of drug repositioning on the Global Health Data Exchange [39] website.

Figure 2 reflects the work methodology used to collect the essential information for this research.

**Fig. 2.**
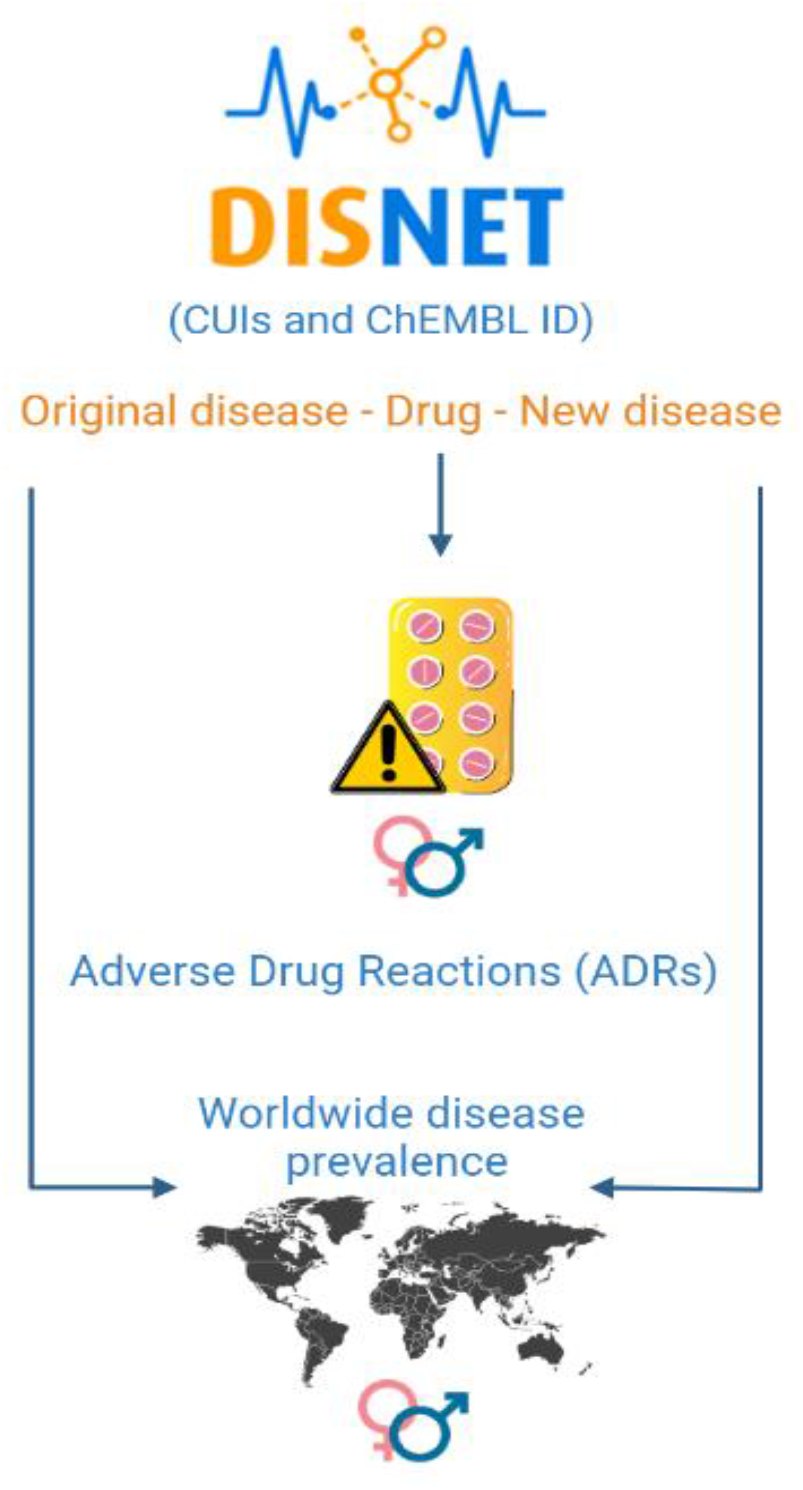
Graphic description of the work methodology carried out in this study.

## IV. RESULTS

### A. Non-rare diseases

The total number of successful drug repositioning cases collected for non-rare diseases from the scientific literature was 136.

The first analysis was carried out to obtain the ADR values of all the drugs involved in the repositioning cases using the VigiAccessTM website, differentiating between the percentage of ADR in men, women and unknown. The number of drug repositioning cases decreased from 136 to 132 because the drugs leading to 4 of the repositioning cases were not present in the VigiAccessTM database.

The mean of ADRs obtained for each of the groups considered was around the following percentages: female (54.28%), male (38.77%), and unknown (6.36%). Based on this first analysis, figure 3 shows how women have a higher percentage of adverse drug reactions in repositioned cases.

**Fig. 3.**
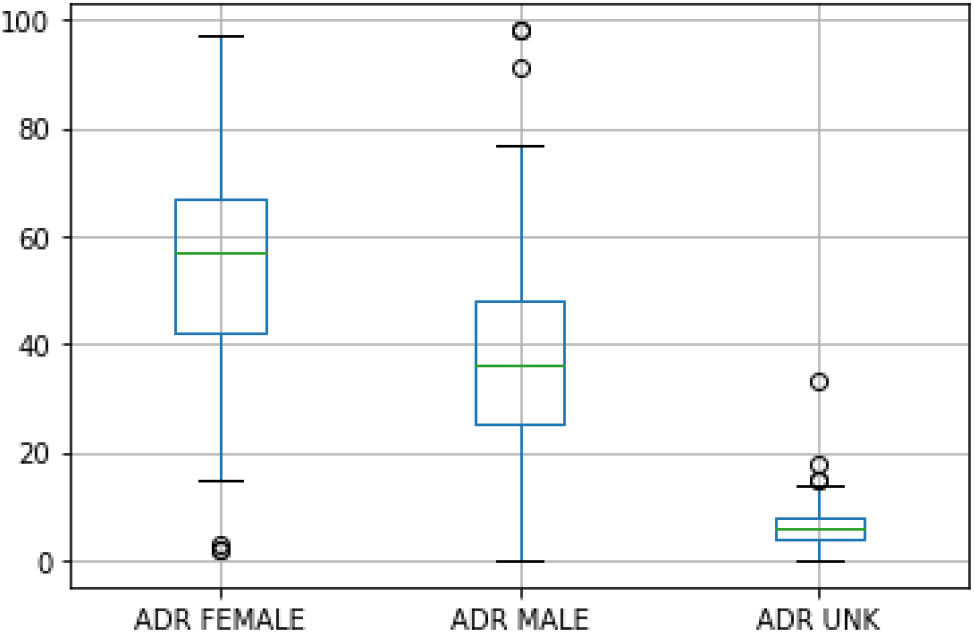
Analysis of the percentage of ADRs presented in women, men and unknown data for all drugs in non-rare diseases.

Then, a Mann-Whitney U Test was performed to compare the ADR data obtained for females, males, and unknowns (code and data available online^1^). Table I shows the results obtained in the cases of repositioning. In it we found that the difference between men and women is statistically significant. The same occurs if we compare men with unknowns or women with unknowns.

**TABLE I.**
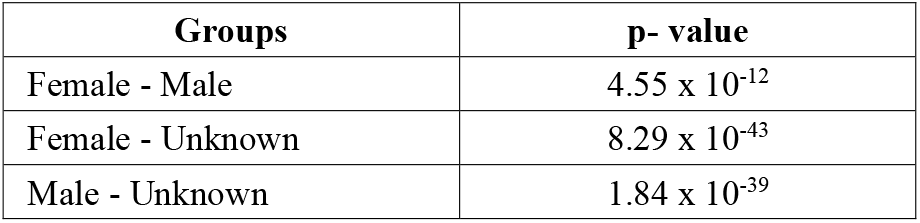
Statistical analysis using Mann-Whitney U Test where males, females and unknown are compared according to their mean ADRS in non-rare diseases

Once we had these results, we moved on to the second part of the study. In this case, out of 136 triplets, we found global prevalence of the original diseases in 100 cases out of 136 and of the new diseases in 92 cases out of 136.

In 33 out of 100 cases of repositioning of original medicines, the overall prevalence was higher in particular gender, while ADRs were higher in the opposite gender. The same results were obtained in 37 out of 92 cases of new medicine repositioning. Thus, we have a total of 70 cases to study. The data have been divided into four groups according to the dominant gender both in the prevalence and in the ADR. The results are shown in Table II..

**TABLE II.** Distribution of ADRS in the drug repositioning cases for non-rare diseases when prevalence gender does not match adr gender.

| Groups | ADR (original disease) | ADR (new disease) | ADR Total |
| --- | --- | --- | --- |
| Prev. male, ADR female | 14 | 18 | 32 |
| Prev. female, ADR male | 16 | 5 | 21 |
| Prev. equal, ADR female | 3 | 6 | 9 |
| Prev. equal, ADR male | 0 | 8 | 8 |

If we combine the results by predominant gender in ADR without accounting for the difference in prevalence, we find that in 41 cases out of 70 females experienced more adverse drug effects even the diseases treated with these drugs are not predominant in the female gender. In the case of men, 29 out of 70 cases show higher adverse drug effects even the diseases treated with these drugs are not male predominant.

### B.Rare diseases

For rare diseases, the total number of successful drug repositioning cases we collected from the scientific literature was 34. All the drugs that form part of the triplets found in the literature are present in the VigiAccessTM database.

Fig. 4 represents the mean ADRs values obtained for each of the groups included in this study in the case of rare diseases. The values corresponding to each of them were: female (49.85%), male (40.41%), and unknown (9.64%). Observations of this first analysis suggest that the number of adverse drug reactions in repositioned ones is higher among females.^1^

**Fig. 4.**
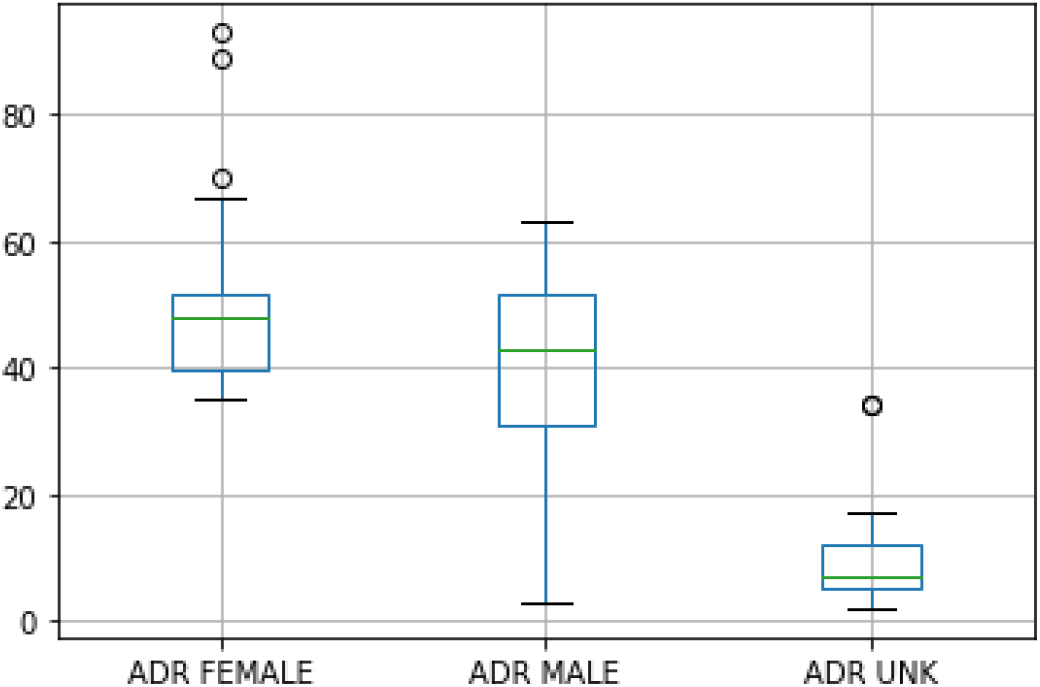
Analysis of the percentage of ADRs presented in women, men and unknown data for all drugs in rare diseases.

As in the previous case, the same statistical methodology was used for the comparison between the different groups studied. Within the repositioning of drugs in rare diseases, it is observed that all the tests that have been carried out comparing the groups with each other are statistically significant (Table III).

**TABLE III.** Statistical analysis using Mann-Whitney U Test where males, females and unknown are compared according to their mean ADRS in rare diseases

| Groups | p-value |
| --- | --- |
| Female - Male | 0.032 |
| Female - Unknown | $5.76 \times 10^{-10}$ |
| Male - Unknown | $6.53 \times 10^{-13}$ |

For the second part of the study regarding prevalence, the results obtained indicate that out of the 34 successful triplets available, we found an overall prevalence of the original diseases in 26 cases out of 34 and of the new diseases in 23 cases out of 34.

Out of 26 the cases of original drug repositioning, in 11 cases the highest number of the global prevalence and the highest number of ADRs did not match in gender. Within the 23 cases of new drug repositioning, in 9 cases the highest number of global prevalence with the highest number of ADRs did not match in gender. Thus, we were left with a total of 20 cases to study. The results are shown in Table IV.

**TABLE IV.** Distribution of ADRS in the drug repositioning cases for rare diseases when prevalence gender does not match ADR gender.

| Groups | ADR (original disease) | ADR (new disease) | ADR Total |
| --- | --- | --- | --- |
| Prev. male, ADR female | 4 | 1 | 5 |
| Prev. female, ADR male | 4 | 8 | 12 |
| Prev. equal, ADR female | 1 | 0 | 1 |
| Prev. equal, ADR male | 2 | 0 | 2 |

In the successful triplets of rare diseases, quite different results are obtained compared to the previous case. Following the same procedure as explained for non-rare diseases, we find that in 6 cases out of 20, females experienced more ADRs even though the diseases treated with these drugs are not predominant in the female gender. In the case of men, this phenomenon is observed in 14 of the 20 cases considered.

## V. DISCUSSION AND CONCLUSIONS

Drug repositioning is a technique that is on the rise because it facilitates the use of existing drugs in new diseases; however, as mentioned throughout the paper, this method has not been studied from a gender perspective.

Numerous studies focus on the importance of drug repositioning in the pharmaceutical industry, as well as a multitude of studies that focus on how men and women react differently to drugs due to biological processes such as pharmacokinetics and pharmacodynamics. But what is really striking about this is that there is no combination of the two fields of study.

Drug repositioning from a gender perspective is an unexplored field of study that has the prospect of being very important in the not-too-distant future.

Our preliminary results show that there is a potentially significant difference between the side effects found in women compared to those observed in men when taking the same drug. These results indicate that in the successful cases of drug repurposing that exist so far, as in *de novo* drug development itself, this action of evaluating a drug for the treatment of a disease is not considering how the ADRs of repositioned drugs affect different to both genders.

According to our findings, for non-rare diseases, adverse effects in women are superior to those in men. However, focusing on rare diseases, we find that it is men who have the higher number of adverse effects. The results obtained in this case are not considered sufficiently significant because there is a very scarce amount of data available.

The limitations of the study include the bounded information obtained in the scientific literature related to successful triplets for rare diseases. The results obtained in this case are not considered sufficiently significant as it is a preliminary analysis. Future research will be necessary for which the number of successful rare disease triplets is not so limited. In the case of non-rare diseases, this limitation also exists, but to a lesser extent.

For this reason, a possible solution to this problem could be to generate synthetic triplets that meet certain criteria to simulate successful drug repurposing triplets in such diseases.

Another limitation was not being able to find the global prevalence of each disease differentiated by sex for all the drug repositioning triplets selected for this study. In the case of the rare disease triplets, this task became even more arduous.

## VI. FUTURE WORKS

The research carried out in this study leaves open many possibilities for future lines of investigation.

On one hand, the relationship between drug repositioning and gender should be explored in more detail More advances and research considering these two points in common could favor results such as those obtained in this work.

On the other hand, within the classification established for drugs, knowing which types of drugs are most likely to cause adverse effects in each gender would be an important advance. With this knowledge, it would be possible to recommend one drug or another to the patient depending on his or her gender. Thus, avoiding possible complications that the patient could develop if he/she had been medicated with the wrong type of drug. This proposal could help in generating a computational mechanism capable of creating personalized medicine based on the most appropriate treatments for the patient according to gender.

Another potential research work could be focused on the withdrawal of drugs from the market. It could be of great interest to know whether the causes of the withdrawal of these drugs are closely related to the adverse effects caused only in one of the genders or whether it is since these drugs produced these effects equally in men and women.

## Acknowledgment

Belen Otero Carrasco’s work is supported by “Formación de Personal Investigador” grant (FPI PRE2019-090912) as part of the project “DISNET (Creation and analysis of disease networks for drug repurposing from heterogeneous data sources applied to rare diseases)” (RTI2018-094576-A-I00) from the Spanish Ministerio de Ciencia, Innovación y Universidades. Lucía Prieto Ssantamaría’s work is supported by “Programa de fomento de la investigación y la innovación (Doctorados Industriales)” from Comunidad de Madrid (grant “IND2019/TIC-17159”). The work is also supported by project “Data-driven drug repositioning applying graph neural networks (3DR-GNN)” (PID2021-122659OB-I00) from the Spanish Ministerio de Ciencia e Innovación.

## Footnotes

1 https://medal.ctb.upm.es/internal/gitlab/b.otero/drug-repositioning-gender

